# A macrophage-smooth muscle cell axis influences vascular remodeling through activation of the EGFR pathway in giant cell arteritis

**DOI:** 10.1101/2024.10.31.621431

**Authors:** Kevin Chevalier, Léa Dionet, Paul Breillat, Margot Poux, Julien Dang, Benoit Terris, Patrick Bruneval, Luc Mouthon, Olivia Lenoir, Pierre-Louis Tharaux, Benjamin Terrier

## Abstract

**Background.:** The role of macrophages and vascular resident cells appears to be predominant in the pathophysiology of giant cell arteritis (GCA). We investigated the role of epithelial growth factor receptor (EGFR) signaling pathway, especially through its activation by heparin-binding epidermal growth factor (HB-EGF) and/or amphiregulin (AREG) in this setting.

**Materials and Methods.:** Serum samples and temporal artery biopsies (TAB) were obtained from patients enrolled in a prospective cohort of systemic vasculitis. Human THP-1, a monocytic cell line, and human aortic vascular smooth muscle cells (VSMC) were used for *in vitro* studies.

**Results.:** Using multiplex immunohistochemistry, TAB from GCA patients showed higher expression of AREG, HB-EGF, EGFR and phospho-EGFR as compared to control arteries. AREG, HB-EGF and EGFR were predominantly expressed by macrophages, whereas EGFR and phosphor-EGFR were expressed by αSMA-positive cells in the media. Increased levels of AREG and HB-EGF were found in culture supernatants of M1 macrophages, whereas M2 macrophages produced only HB-EFG. AREG and HB-EGF did not increase the production of pro-inflammatory cytokines by THP-1 or macrophages but activated the p38 MAPK pathway. Using transcriptomic and Western blot analysis of human aortic VSMC, AREG and especially HB-EGF induced cell proliferation pathway, enhanced interferon alpha and gamma responses, and activation of the MAPK pathway. Finally, AREG and HB-EGF increased both VSMC proliferation and migration, which were completely inhibited by AG1478, an EGFR inhibitor.

**Conclusion.:** We show that both AREG and HB-EGF may play a role in the pathophysiology of GCA, especially in the remodeling phase of the disease.

## Introduction

Giant cell arteritis (GCA) is a granulomatous/giant cell large-vessel vasculitis that primarily affects the cephalic arteries and aorta (1,2). It is the most common vasculitis among people over 50 years of age, with a peak incidence at 70-75 years of age and a female predominance (1,2). The pathophysiology of GCA is commonly described as a 4-phase model (3,4): i) tolerance breakdown and activation of resident dendritic cells in the arterial adventitia (5), ii) CD4^+^ T cells recruitment with Th1 and Th17 polarization (6,7), iii) recruitment of CD8^+^ T cells and monocytes into the vessels and differentiation into macrophages with pro-inflammatory properties (8,9), and iv) vascular remodeling leading to ischemic complications, the latter being mainly dependent on vascular smooth muscle cells (VSMC) and myofibroblasts invasion of the media and the intima (10). The role of monocytes and macrophages, VSMC and myofibroblasts appears to be prominent in the pathophysiology of GCA (10). Currently, the standard-of-care for GCA is based on glucocorticoids (GCs), which are remarkably effective. However, approximately 15% of patients experience permanent visual impairment despite treatment and 40 to 50% of patients with GCA experience relapses when GCs are tapered (11,12). Treatment is also unable to prevent large vessel damage leading to aortic dilatation (13). Finally, GCs-related adverse events are common in this elderly population (12). New therapeutic targets have been identified to reduce exposure to GCs, including the blockade of the interleukin (IL)-6 receptor or the IL-17. However, these therapies are costly, further weaken the immune system, and may have a limited effect on VSMC and myofibroblasts proliferation and vascular remodeling (2,3,14). Therefore, elucidation of the pathophysiological mechanisms of vascular remodeling and identification of novel strategies that do not target the immune system may be of interest in the treatment of patients with GCA.

The epidermal growth factor receptor (EGFR) signaling pathway is critically involved in a variety of cellular processes, including cell cycle, growth and differentiation (15). EGFR-blocking agents have been established as treatments for several human malignancies. However, recent studies have shown that targeting the EGFR pathway may be effective beyond cancer (16). More specifically, the role of heparin-binding epidermal growth factor (HB-EGF) and amphiregulin (AREG), two major ligands of EGFR, in inflammatory diseases and tissue fibrosis in both animal models and humans has emerged in the literature (17–21). We previously identified AREG as a pro-inflammatory mediator in a murine model of glomerulonephritis, leading to increased renal infiltration of myeloid cells and having direct effects on monocyte/macrophage chemotaxis, polarization, proliferation, survival and cytokine secretion (21). In addition, we identified HB-EGF as an autocrine mediator that induces a phenotypic switch of podocytes in rapidly progressive glomerulonephritis and proliferation of intrinsic glomerular cells (17). Thus, the EGFR signaling pathway might be involved in vascular inflammatory diseases such as GCA.

In the present study, we investigated the role of the EGFR signaling pathway in the pathophysiology of GCA and potentially identified novel therapeutic targets.

## Material and methods

### Patients

Patients with GCA were defined according to the 2022 ACR/EULAR criteria (22). Controls were healthy controls for serum analysis and patients in whom GCA was suspected but the diagnosis was finally excluded and who had no other inflammatory disease for temporal artery biopsy (TAB) analysis. Thus, we collected and used for this study 21 serum samples from GCA patients, 15 from healthy controls, 7 TAB from GCA patients, and 8 TAB from control patients. Serum samples and TAB were obtained from patients enrolled in the VASCO (VASculitis COhort) study, a prospective cohort of patients with systemic vasculitides (NCT04413331), which includes a biological collection approved by the Direction de la Recherche Clinique et Innovation (DRCI) and the Ministère de la Recherche (N°2019-3677).

### Temporal artery biopsy specimens

TAB were obtained from the pathology departments of the Cochin Hospital and the Georges Pompidou European Hospital, Paris, France. Biopsies were fixed in formalin and embedded in paraffin, and 4-μm-thick sections were processed for histopathology or immunohistochemistry.

### Immunohistochemistry

Paraffin-embedded sections were deparaffinized and rehydrated through Xylene and ethanol baths. Antigen retrieval was performed after the slides were placed in citrate buffer (pH 6) at 95°C in a retrieval chamber (Decloacking ChamberTM NxGen, Biocare Medical, Pacheco, USA) for 25 minutes and then cooled on ice for 30 minutes. The slides were then permeabilized with 0.1% TBS-Triton solution for 10 minutes and incubated in 3% TBS-T-BSA for 30 minutes. The slides were then incubated with the appropriate primary antibodies overnight at +4°C (see details in the **Supplementary material**). Slides were washed in TBS-T and then incubated with peroxidase-conjugated antibodies (Histofine®, Nichirei Biosciences, Japan) for 2 hours at room temperature. After washing in TBS-T, staining was revealed with 3,3’-diaminobenzidine (DAB) for 2-6 minutes and counterstained with hematoxylin. Sections were scanned with Olympus slideview vs200 (Tokyo, Japan) and processed with QuPath software (23).

### Multiparametric immunofluorescence staining

Multiparametric immunofluorescence staining based on tyramide signal amplification (OPAL, Akoya, #OP7DS2001KT) was used to stain TAB sections with a set of different markers. Briefly, slides were deparaffinized as described above, incubated with blocking antibody diluent (Akoya, ARD1001EA) for 10 minutes, and then incubated with primary antibodies overnight at +4°C (see details in the **Supplementary material**). The slides were then washed in TBS-T, incubated with Opal Polymer HRP (anti-mouse and anti-rabbit, Akoya, ARH1001EA) for 10 minutes and labeled with Opal dyes (Akoya, OP-01000, OP-001001, OP-001003, OP-001004, OP-001006, OP-001007, 1:100). The slides were then heated in citrate buffer (pH 6) in a microwave oven for 10 minutes and cooled on ice for 30 minutes. The above sequence was repeated for the whole panel of primary antibodies. The images were analyzed and quantified using HALO software (Indica Labs, Albuquerque, USA).

### Cell lines and cell culture

Three cell lines were used in this study: wild-type THP-1 cells (Merck, 88081201-1VL), a human monocyte cell line derived from a patient with acute monocytic leukemia, THP1-Blue^TM^ which are NF-κB reporter monocytes (Invivogen, thp-nfkb), and HAoSMC (Human Aorta, Smooth Muscle Cells; Promocell, C-12533). THP1-Blue^TM^ were specifically designed for monitoring the NF-κB signal transduction pathway in a physiologically relevant cell line. Both cell lines were grown in a humid incubator at 5% CO2, 37°C in the appropriate culture media: Smooth Muscle Cell Growth Medium 2 (Promocell, C-22062) and SupplementMix (Promocell, C-39267) with 1% penicillin/streptomycin (P/S) for HAoSMCs and RPMI 1640 1X + glutamax + 25mM Hepes (Gibco, 72400-021) with 10% fetal bovine serum (FBS) and 1% P/S for THP-1. The cultures were maintained at a concentration of 200,000 cells/mL for HAoSMC and 1,000,000 cells/mL for THP-1. These concentrations were used throughout the experiment.

### Stimulation of HAoSMC and THP-1 cells

Cell lines were stimulated with: lipopolysaccharide (LPS) 100 ng/mL (Sigma Aldrich, L2630-10MG), tumor necrosis factor-α (TNF-α) 50 ng/mL (R&D, 210-TA-005), AREG 50 ng/mL (R&D, 262-AR-100) and HB-EGF 50 ng/mL (Peprotech, 100-47). Cells were also inhibited with an EGFR inhibitor, AG1478, at 500 nM (Calbiochem, 658548).

### Measurement of NF-κB activity in THP1-cells by QUANTI-Blue

Briefly, we incubate THP1-Blue cells with 180 μL of QUANTI-Blue™ Solution (Invivogen, rep-qbs) and 20 μL of supernatant containing different stimuli as described above in a 96-well plate. After incubation at 37°C for 30 minutes, we measure optical density (O.D) at 620-655 nm using a microplate reader (Spectrostar Nano, BMG Labtech).

### Differentiation and polarization of THP-1 cell line

THP-1 monocytic cells were differentiated into macrophages after 24 hours of stimulation with phorbol 12-myristate 13-acetate (PMA) 100 ng/mL (Sigma Aldrich, P8139-1MG), then polarized into M1 or M2 macrophages after 48 hours of culture with 20 ng/mL interferon-γ (INF-γ) (R&D, 285-IF-100) and 1000 pg/mL LPS (Sigma Aldrich, L2630-10MG) for M1 polarization, and 20 ng/mL IL-4 (Peprotech, 200-04-20UG) and 20 ng/mL IL-13 (Peprotech 200-13A-10UG) for M2 polarization.

### Enzyme-Linked Immunosorbent Assay (ELISA) and Luminex

Measurement of AREG and HB-EGF in patient serum was performed by ELISA according to the manufacturer’s instructions (Human Amphiregulin Quantikine ELISA Kit, R&D Systems, DAR00 and Human HB-EGF DuoSet ELISA, R&D Systems, DY259B, respectively). Cytokine secretion by THP-1 cells, M1 and M2 macrophages was measured using Luminex® (Human Magnetic Luminex® Assays Premixed Multiplex, R&D Systems, LXSAHM-07) according to the manufacturer’s instructions.

### Western blot

Cell lysates were prepared with radioimmunoprecipitation lysis buffer (RIPA) (150 mM NaCl, 50 mM Tris-HCl, 2 mM EDTA, 0.5% sodium deoxycholate, 0.2% sodium dodecyl sulfate (SDS), and 1% NP40 in MilliQ) containing protease inhibitors (Roche, 11836170001) and phosphatase inhibitors (Roche, 4906845001). Equal amounts of protein were loaded onto pre-cast polyacrylamide electrophoresis gels (NuPAGE™ 4 to 12%, Bis-Tris, 1.0 mm, Midi Protein Gels, Thermo Fisher, WG1401BOX) in NuPAGE SDS MES or NuPAGE SDS MOPS migration buffer (Thermo Fisher, NP000202 and NP000102) for separation and transfer to PVDF (polyvinylidene difluoride) membranes (Thermo Fisher, IB24001). The membranes were then blocked in 5% TBS-T-milk mixture and incubated with different primary antibodies for 4 hours (see details in the **Supplementary material**). After washes in TBS-T, the membranes were incubated with various secondary antibodies (see details in the **Supplementary material**). Bands were detected by chemiluminescence (Clarity Western ECL substrate, Bio-Rad, 1705061). Amersham ImageQuant™ 800 device (Grosseron, Couëron, France) was used to visualize the bands, and densitometric analysis was used for quantification using ImageJ software (NIH, USA). All bands were normalized in housekeeping protein (tubuline or GAPDH).

### Cell migration assay

HAoSMC were cultured with standard culture medium in Culture-Insert 2-well plates (Ibidi, 80242) for 24 hours to reach quiescence. The cells were washed with PBS and starved with smooth muscle cell growth medium 2 containing 1% FBS + 1% insulin-transferrin-selenium (Gibco, 41400045) + 1% P/S overnight. After further washing with PBS, the inserts were removed and the cells were incubated with LPS (100 ng/mL), TNFα (50 ng/mL), AREG (50 ng/mL), HB-EGF (50 ng/mL), or AG1478 (500 nM) for 72 hours in smooth muscle cell growth medium 2 containing 1% FBS + 1% insulin-transferrin-selenium (Gibco, 41400045) + 1% P/S. The cells were incubated into the inserts in a stage top chamber (OKOLAB, H301-T-UNIT-BL-PLUS) at 5% CO2, 37°C. Images of cells migrating into the gap were taken at 0 hours and then every hour until the gap was closed thanks to an inverted microscope (Eclipse Ti, Nikon) combined with Nikon’s NIS-Elements imaging software (Nikon, Melville, USA). Images were compared to quantify the rate of cell migration using also the NIS-Elements imaging software (Nikon, Melville, USA).

### Statistics

Data are expressed as mean ± SEM. Statistical analyses were performed using GraphPad Prism software (GraphPad Software, La Jolla, CA). Comparisons between 2 groups were made using t-tests. Comparisons between multiple groups were made by ANOVA test. A p-value <0.05 was considered significant.

## Results

### EGFR and its ligands are expressed in temporal arteries from GCA patients

To evaluate the potential role of EGFR ligands (AREG, HB-EGF, epithelial growth factor (EGF), epiregulin (EREG), tumor growth factor α (TGF-α), and betacellulin) and the EGFR signaling pathway (EGFR and phospho-EGFR) in the pathogenesis of GCA, we first performed immunohistochemistry in TAB from GCA patients and controls. All EGFR ligands, including AREG, HB-EGF, EGF, EREG, TGF-α and betacellulin, were highly expressed in the media of TAB from GCA patients but not from control arteries (**Figure 1A** and **Supplementary Figure 1**). To confirm the activation of the EGFR pathway, we stained for EGFR and pEGFR and we observed higher expression of EGFR and pEGFR in the media of TAB from GCA patients compared to controls (**Figure 1A**).

**Figure 1.**
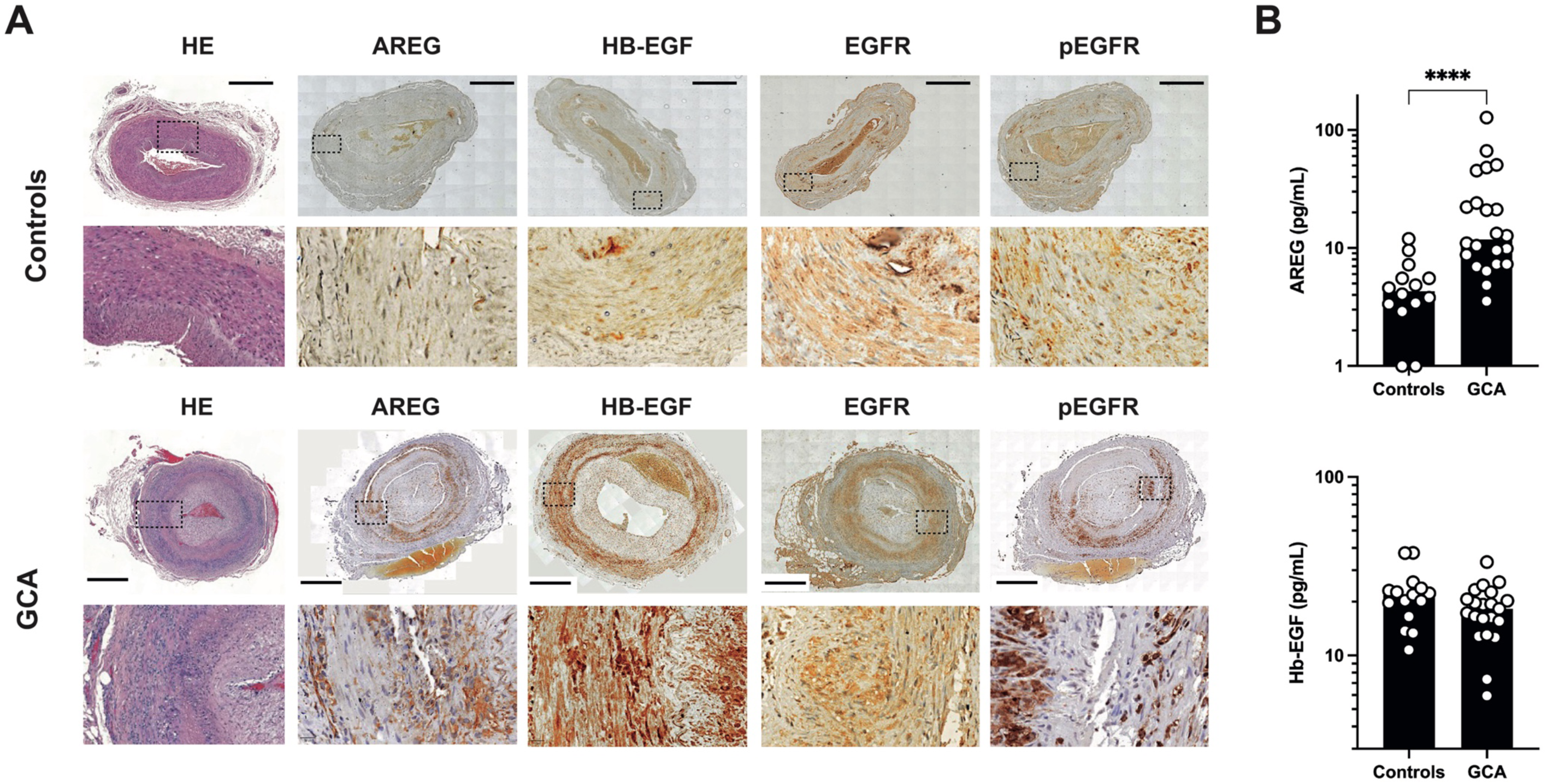
Expression of AREG, HB-EGF, EGFR and pEGFR in temporal arteries from GCA and controls. **(A)** Hematoxylin-eosin staining of temporal artery biopsy from a control and a GCA patient and immunohistochemical staining of AREG, HB-EGF, EGFR and pEGFR of the same temporal arteries. Illustrative pictures show increased expression of AREG, HB-EGF, EGFR and pEGFR in the media from temporal arteries from GCA patient (Magnification x 2 and x 20, scale bar 500 μm). **(B)** Serum AREG and HB-EGF from controls and GCA patients. Each dot represents a single patient. P values were determined by the two-sided Kruskal-Wallis test, followed by Dunn’s post test for multiple group comparisons. *P <0.05; **P <0.01; ***P <0.001, ****P <0.0001.

Since AREG and HB-EGF are well described as pro-inflammatory ligands, we decided to focus on these two ligands. In addition to tissue analysis, we measured serum levels of AREG and HB-EGF. Serum AREG levels were significantly higher in GCA than in controls, whereas HB-EGF levels were similar between GCA and controls (**Figure 1B**).

We next evaluated cell types expressing AREG, HB-EGF, EGFR and pEGFR using multiparametric immunofluorescence staining on TAB. Quantification of AREG, HB-EGF, EGFR and pEGFR-expressing cells confirmed higher expression in GCA compared to controls (**Figure 2A**). We analyzed the colocalization of AREG, HB-EGF, EGFR and pEGFR, with macrophages defined as CD68-positive cells and vascular resident cells, specifically VSMC, commonly found in the media, and myofibroblasts, commonly found in the intima, both defined as α-smooth muscle actin (α-SMA)-positive cells. We observed a mild expression of AREG and HB-EGF in the adventitia of the temporal artery from controls, and an expression of EFGR, but not pEGFR, by some α-SMA-positive cells in the media resembling VSMC (**Figure 2B**). In contrast, analysis of TAB from GCA patients showed strong expression of AREG and HB-EGF by macrophages invading the media, with very few α-SMA-positive cells expressing HB-EGF but not AREG (**Figure 2C**). Both infiltrating macrophages and α-SMA-positive cells in the media suggestive of VSMC strongly expressed EGFR and pEGFR in GCA patients, with some α-SMA-positive cells in the hyperplasic intima, probably myofibroblasts, also expressing pEGFR (**Figure 2C**). Collectively, these findings support a role for the EGFR signaling pathway in the pathophysiology of GCA.

**Figure 2.**
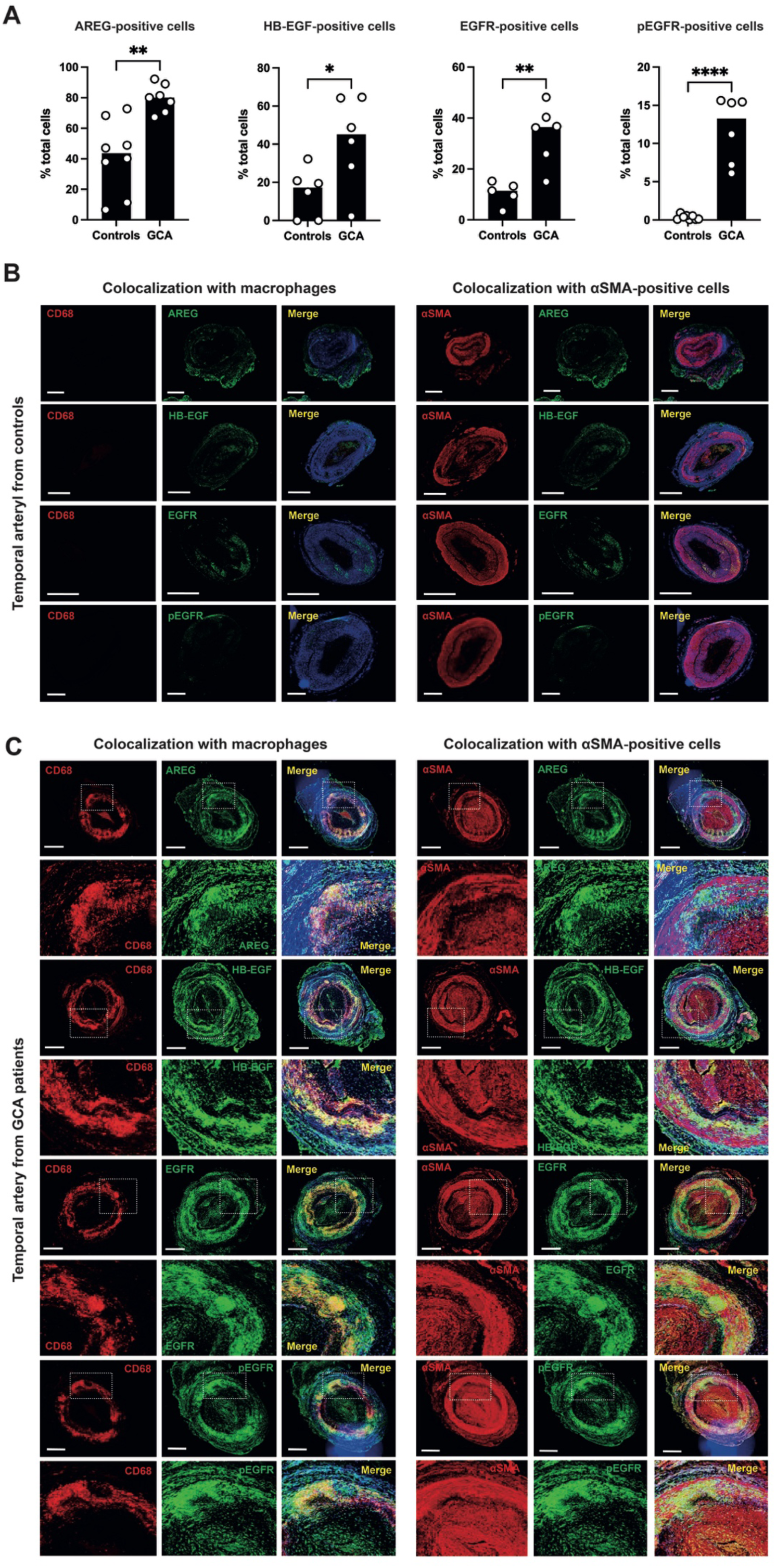
Characterization of cell types expressing AREG, HB-EGF, EGFR and pEGFR within temporal arteries from GCA patients and controls. **(A)** Quantification of AREG, HB-EGF, EGFR and pEGFR-positive cells among temporal arteries from controls and GCA patients. Data show an increased expression of AREG, HB-EGF, EGFR and pEGFR in GCA compared to controls. P values were determined by the two-sided Kruskal-Wallis test, followed by Dunn’s post test for multiple group comparisons. *P <0.05; **P <0.01; ***P <0.001, ****P <0.0001. **(B)** Opal multiplex immunohistochemistry of temporal artery from a control using AREG, HB-EGF, EGFR, pEGFR, CD68 and αSMA antibodies. Illustrative pictures show a mild expression of AREG and HB-EGF in the adventitia of the temporal artery from controls, and the expression of EFGR but not pEGFR by αSMA-expressing cells in the media (Magnification x 2, scale bar 500 μm). **(C)** Opal multiplex immunohistochemistry of temporal artery biopsy from a GCA patient using AREG, HB-EGF, EGFR, pEGFR, CD68 and αSMA antibodies. Illustrative pictures show a strong expression of AREG and HB-EGF by macrophages invading the media, with very few αSMA-positive cells expressing HB-EGF but not AREG, and both infiltrating macrophages and αSMA-positive cells in the media strongly expressed EGFR and pEGFR (Magnification x 2 and x 20, scale bar 500 μm).

### AREG and HB-EGF are expressed by M1 and M2 macrophages and activate the P38 MAPK pathway

We then analyzed the expression and function of AREG and HB-EGF by monocytes and/or macrophages, using the THP-1 monocytic cell line and THP-1-derived macrophages (i.e., M1 and M2 macrophages).

We used cytospin and ibidi slides to visualize AREG and HB-EGF expression of THP-1 and THP-1-derived macrophages, respectively, by confocal microscopy. AREG was expressed by M1 and M2 macrophages but not by THP-1 cells, with stronger expression in M1 macrophages, whereas HB-EGF was expressed in both THP-1 cells and THP-1-derived M1 and M2 macrophages (**Figure 3A**). To assess the secretion of AREG and HB-EGF by monocytes and macrophages, we next measured AREG and HB-EGF levels in the culture supernatants of THP-1 and THP-1-derived M1 and M2 macrophages. M1 macrophages secreted both AREG and HB-EGF, whereas M2 macrophages secreted only HB-EGF. In contrast, THP-1 cells did not secrete AREG or HB-EGF (**Figure 3B**). The upregulation of *AREG* and *HB-EGF* in THP-1-derived macrophages was also confirmed at the transcriptomic level (**Figure 3C**).

**Figure 3.**
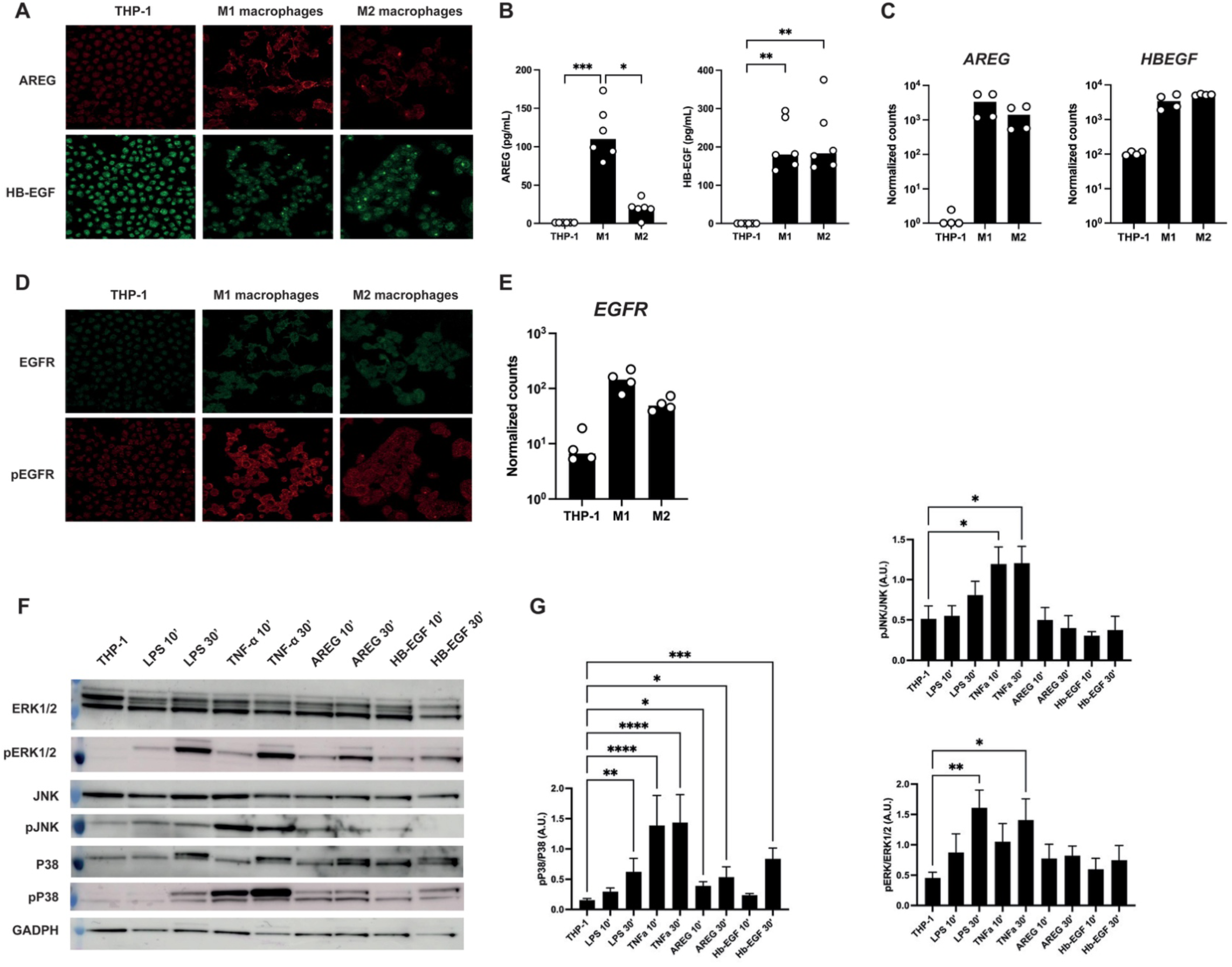
Expression of AREG, HB-EGF, EGFR and pEGFR by THP-1 and THP-1-derived macrophages and stimulation of the MAPK pathway in THP-1 cells by AREG and HB-EGF. **(A)** Confocal microscopy of THP-1 and THP-1-derived M1 and M2 macrophages using AREG and HB-EGF antibodies. Illustrative pictures show increased expression of AREG and HB-EGF in THP-1-derived M1 and M2 macrophages. **(B)** Quantification by ELISA of AREG and HB-EGF in culture supernatants of THP-1 and THP-1-derived M1 and M2 macrophages, showing a significantly increased expression of AREG in THP-1-derived M1 macrophages only and HB-EGF in both THP-1-derived M1 and M2 macrophages, compared to THP-1. **(C)** Transcriptomic analysis of THP-1 and THP-1-derived M1 and M2 macrophages, showing a significantly increased expression of AREG and HB-EGF in both THP-1-derived M1 and M2 macrophages, compared to THP-1. **(D)** Confocal microscopy of THP-1 and THP-1-derived M1 and M2 macrophages using EGFR and pEGFR antibodies. Illustrative pictures show increased expression of EGFR in THP-1-derived M1 and M2 macrophages, and pEGFR mainly in THP-1-derived M1 macrophages. **(E)** Transcriptomic analysis of THP-1 and THP-1-derived M1 and M2 macrophages, showing an increased expression of EGFR in both THP-1-derived M1 and M2 macrophages, mainly M1 macrophages, compared to THP-1. (**F**) Illustrative Western blot of THP-1 cells after 10- and 30-minutes stimulation by LPS, TNFα, AREG or HB-EGF. (**G**) Quantification of Western blot measuring p38, JNK and ERK phosphorylation after stimulation. Both AREG and HB-EGF induced a significant increase in p38 phosphorylation compared to no stimulation. P values were determined by the two-sided Kruskal-Wallis test, followed by Dunn’s post test for multiple group comparisons. *P <0.05; **P <0.01; ***P <0.001, ****P <0.0001.

We also visualize EGFR and pEGFR expression of THP-1 and THP-1-derived M1 and M2 macrophages. EGFR and pEGFR were weakly expressed in the nucleus of THP-1 cells, while THP-1-derived macrophages expressed EGFR and pEGFR in their cytoplasm, mainly M1 macrophages (**Figure 3D**). The upregulation of *EGFR* was also higher in THP-1-derived macrophages at the transcriptomic level, especially in M1 macrophages (**Figure 3E**). We evaluated the ability of AREG and HB-EGF to stimulate the MAPK pathway in THP-1 cells 10 and 30 minutes after stimulation with LPS, TNF-α, AREG and HB-EGF by Western blot. Both AREG and HB-EGF significantly induced P38 phosphorylation compared to no stimulation, supporting a role for AREG and HB-EGF in some biological processes (**Figure 3F and 3G**).

Since macrophages express both AREG and HB-EGF as well as EGFR, we sought to determine whether AREG and HB-EGF could have an autocrine or paracrine effect on THP-1 cells or THP-1-derived macrophages. We first used the THP1-Blue™ reporter cell line to measure the effect of AREG and HB-EGF on NFκB activation. AREG and HB-EGF did not increase NFκB activation in either THP-1 or THP-1-derived macrophages (**Figure 4A**). We also evaluated the effect of AREG and HB-EGF on the production of inflammatory cytokines by THP-1 cells and THP-1-derived M1 and M2 macrophages and did not observe any effect of AREG and HB-EGF on the secretion of IL-1β, IL-6, IL-8, TNF-α or IFN-γ (**Figure 4B and 4C**).

**Figure 4.**
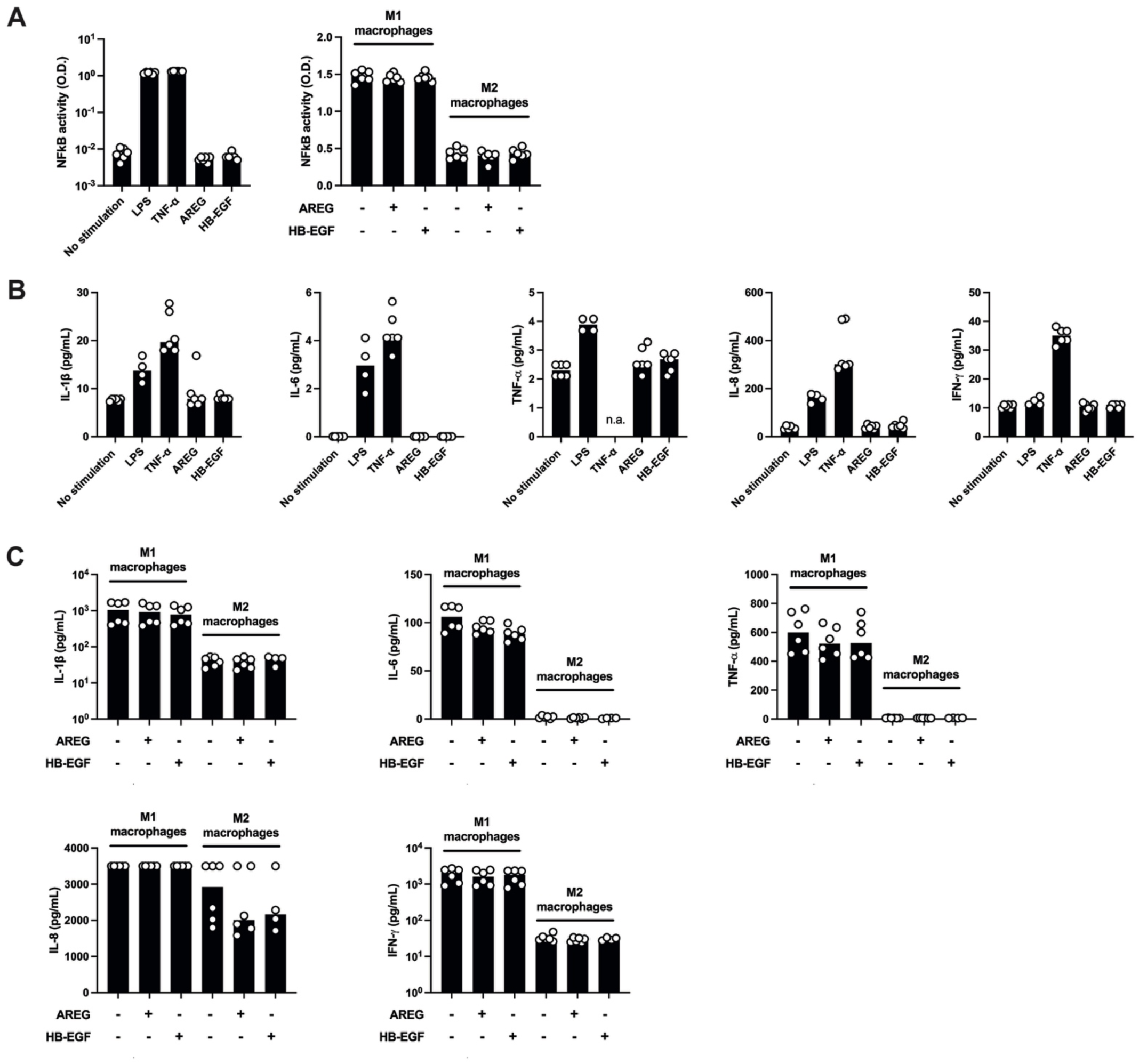
Absence of proinflammatory effect of AREG and HB-EGF on THP-1 cells and THP-1-derived macrophages. **(A)** Measurement of NF-κB activity using QUANTI-Blue in THP-1 and THP-1-derived M1 and M2 macrophages after stimulation with LPS, TNFα, AREG or HB-EGF. NF-κB activity is increased by LPS and TNFα in THP-1 cells but not by AREG or HB-EGF stimulation. NF-κB activity is higher in THP-1-derived M1 macrophages than in THP-1-derived M2 macrophages, without any effect of AREG or HB-EGF stimulation. **(B)** Measurement of pro-inflammatory cytokines by Luminex® in culture supernatants of THP-1 cells after stimulation with LPS, TNFα, AREG or HB-EGF. Both LPS and TNF-α increased the secretion of pro-inflammatory cytokines, but not AREG or HB-EGF. **(C)** Measurement of pro-inflammatory cytokines by Luminex® in culture supernatants of THP-1-derived M1 and M2 macrophages after stimulation with AREG or HB-EGF. Pro-inflammatory cytokines were higher in culture supernatants of THP-1-derived M1 macrophages than of M2 macrophages, but without any effect of AREG or HB-EGF stimulation.

Taken together, these results support that macrophages express and secrete both AREG and HB-EGF, with AREG secreted primarily by M1 macrophages and HB-EGF secreted by both M1 and M2 macrophages, and that both activate the P38 MAPK pathway in monocytes.

### AREG and HB-EGF induce activation of the MAPK pathway and proliferation and migration of VSMC

Because α-SMA-positive cells in the media, mainly VSMC, express both EGFR and its phosphorylated form, pEGFR, we examined the effect of AREG and HB-EGF stimulation on VSMC.

First, we performed transcriptomic analysis of human aortic VSMC stimulated or not with AREG or HB-EGF. Differential gene expression and gene set enrichment analysis (GSEA) revealed enrichment in proliferation, oxidative phosphorylation and interferon alpha and gamma response signatures in HB-EGF- and AREG-stimulated VSMC (**Figure 5A)**.

**Figure 5.**
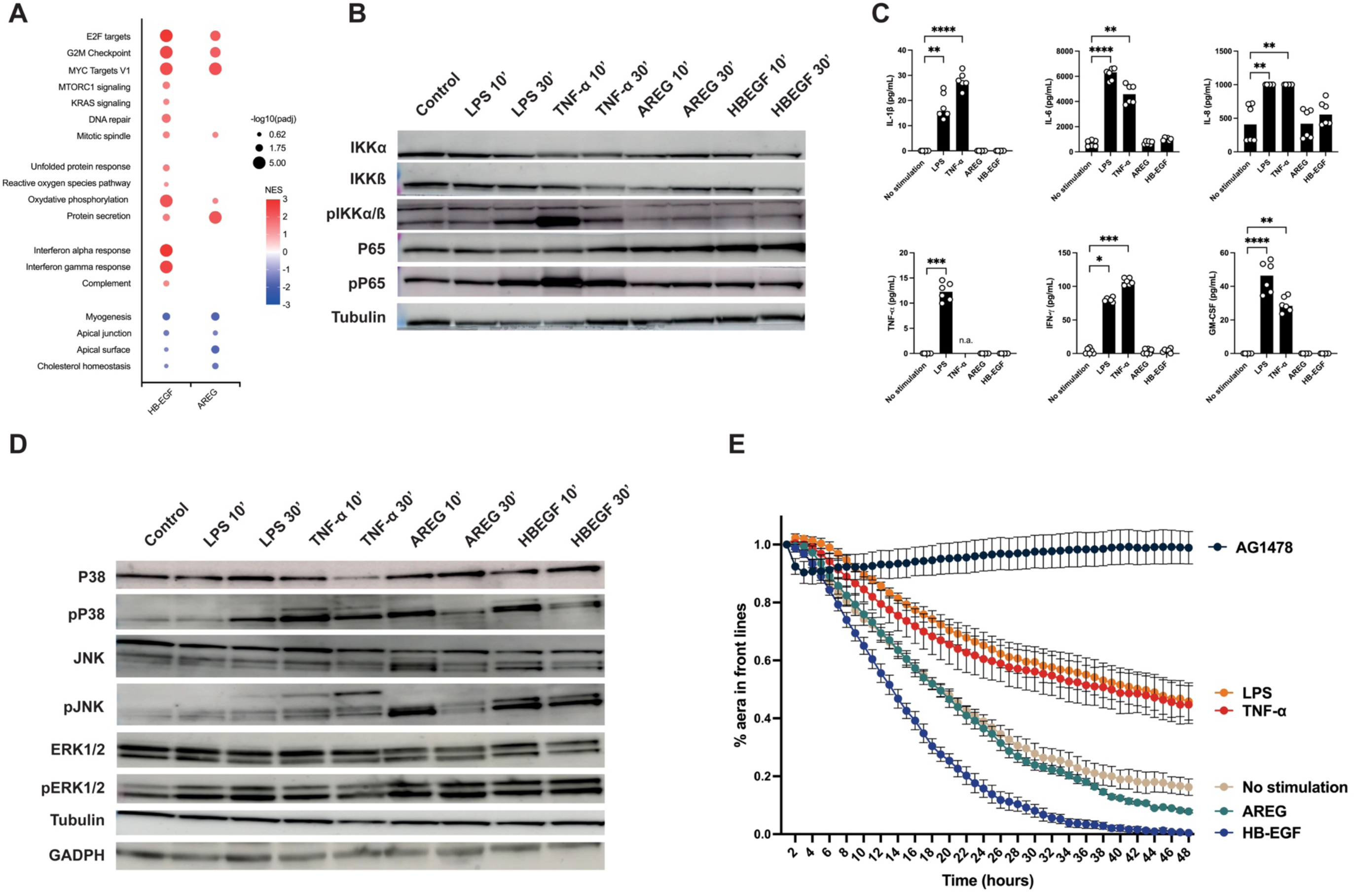
Effects of AREG and HB-EGF on human aortic vascular smooth muscle cells. (**A**) Transcriptomic analysis and gene set enrichment analysis of human aortic VSMC treated or not with AREG or HB-EGF. GSEA analysis revealed enrichment in proliferation, oxidative phosphorylation and interferon alpha and gamma response signatures in HB-EGF- and AREG-stimulated VSMC. (**B**) Illustrative Western blot of human aortic VSMC after 10- and 30-minutes stimulation by LPS, TNFα, AREG or HB-EGF, showing that AREG and HB-EGF do not stimulate the NFκB pathway. **(C)** Measurement of pro-inflammatory cytokines by Luminex® in culture supernatants of human aortic VSMC after stimulation with LPS, TNFα, AREG or HB-EGF. Both LPS and TNF-α increased the secretion of pro-inflammatory cytokines, but not AREG or HB-EGF. (**D**) Illustrative Western blot of human aortic VSMC after 10- and 30-minutes stimulation by LPS, TNFα, AREG or HB-EGF, showing increased phosphorylation of P38, JNK and ERK induced by AREG or HB-EGF. **(E)** Live cell imaging assessing proliferation and migration after treatment with LPS, TNF-α, AREG, HB-EGF, and AG1478, an EGFR inhibitor. AREG and especially HB-EGF increased VSMC proliferation and migration, whereas AG1478, an EGFR inhibitor, completely inhibited VSMC proliferation and migration.

We evaluated the ability of AREG and HB-EGF to stimulate the NFκB in human aortic VSMC at 10 minutes and 30 minutes with LPS, TNF-α, AREG and HB-EGF. In contrast to TNF-α, AREG and HB-EGF did not stimulate the NFκB pathway (**Figure 5B**). LPS and TNF-α induced the production of inflammatory cytokines in the culture supernatants of human aortic VSMC, whereas AREG and HB-EGF did not (**Figure 5C**).

We next evaluated the ability of AREG and HB-EGF to stimulate the MAPK pathway in human aortic VSMC. Both AREG and HB-EGF significantly induced ERK, P38 and JNK phosphorylation compared with no stimulation (**Figure 5D**). Based on these results, we hypothesized that the stimulation of the MAPK pathway in VSMC could increase both proliferation and migration. Using live cell imaging, we showed that AREG and especially HB-EGF increased VSMC proliferation and migration, whereas AG1478, an EGFR inhibitor, completely inhibited VSMC proliferation and migration (**Figure 5E**), suggesting a key role for the EGFR pathway in VSMC proliferation and migration and vascular remodeling in GCA, particularly through activation of the MAPK pathway.

## Discussion

We aimed to investigate the previously unexplored role of the EGFR pathway in GCA. The main findings of our study were: (i) AREG, HB-EGF, EGFR and pEGFR are highly expressed in TAB from GCA patients, (ii) AREG and HB-EGF are highly expressed by macrophages infiltrating TAB from GCA, while EGFR and pEGFR are expressed by VSMC, (iii) both AREG and HB-EGF induce VSMC proliferation and migration in part by stimulating the MAPK pathway, and proliferation and migration is completely inhibited by EGFR inhibitor. Our findings suggest that EGFR ligands may be secreted by macrophages invading the vessel wall in GCA and may affect macrophages and vascular resident cells, both expressing EGFR (**Figure 6**).

**Figure 6.**
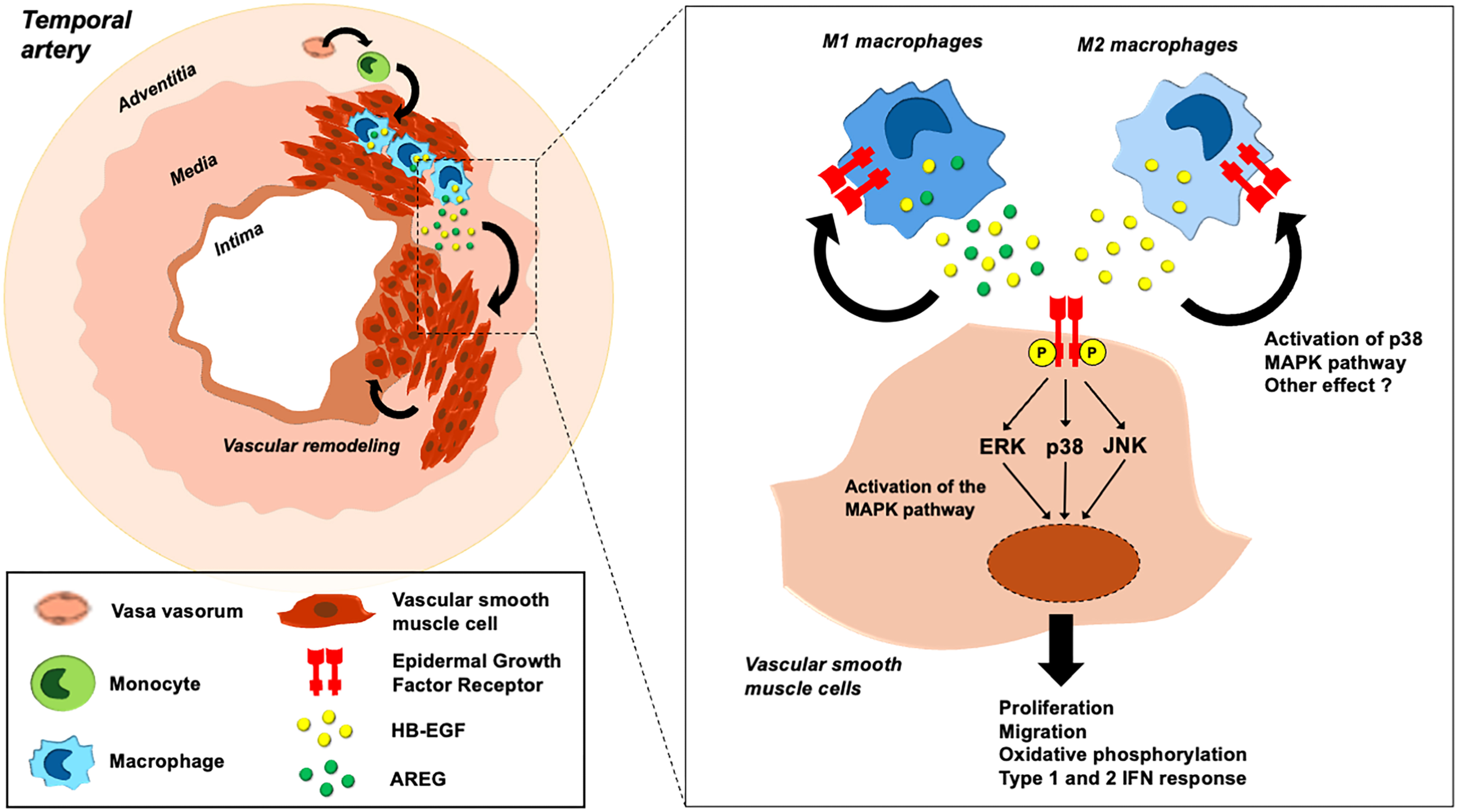
Role of the EGFR signaling pathway in GCA. In temporal artery biopsies from patients with giant cell arteritis, macrophages invading the vessel wall secreted EGFR ligands (i.e., amphiregulin (AREG) and heparin-binding EGF (HB-EGF)). M1 macrophages secreted both AREG and HB-EGF, whereas M2 macrophages secreted only HB-EGF. Both ligands can affect both macrophages and vascular resident cells, both of which express EGFR. In monocytes/macrophages, AREG and HB-EGF induce activation of the p38 MAPK pathway and may have other effects. In vascular smooth muscle cells, AREG and HB-EGF induce activation of the MAPK pathway through phosphorylation of ERK, p38, and JNK, resulting in increased proliferation, migration, oxidative phosphorylation, and type 1 and type 2 interferon (IFN) response. These mechanisms can lead to vascular remodeling of the arterial wall, resulting in ischemic clinical signs.

AREG has been shown to play several roles in physiology, first by promoting pro-inflammatory responses with the production of pro-inflammatory cytokines such as IL-6 or GM-CSF by epithelial cells or synoviocytes (19,20,24), and second by playing a role in type 2 inflammation and tissue repair and regeneration (24). Therefore, AREG can be expressed by multiple cell populations in a variety of inflammatory conditions (25), such as peripheral blood mononuclear cells (20), T helper 2 cells, innate lymphoid cells type 2, regulatory T cells, mucosal-associated invariant T cells (26), dendritic cells (27), mast cells (28), eosinophil cells (29) or macrophages (30,31). Previous data showed that AREG is significantly expressed by M1 macrophages compared to M2 or non-polarized peritoneal or alveolar macrophages in mice and that AREG production temporally paralleled the production of classical M1 biomarkers (i.e. TNF-α, IL-1β or IL-6) (24), suggesting that AREG could be a biomarker of M1 macrophages. The production of AREG in macrophages appears to be dependent on TLR4 signaling, similar to the polarization of macrophage into M1 macrophages, but also through MAPK signaling (24). We observed that LPS and IFN-γ increased AREG production by M1 macrophages, but also AREG could activate the MAPK pathway in monocytes, suggesting an amplification loop in AREG production by monocytes/macrophages. IFN-γ is also highly produced by Th1 cells in the GCA, which could be responsible for both activation of VSMC and polarization of macrophages into M1 macrophages with increased production of AREG (3,8). Expression of HB-EGF by immune cells, mainly CD4^+^ T cells (32) or macrophages (33,34), have also been demonstrated in several inflammatory conditions, such as rapidly progressive glomerulonephritis (17), lupus nephritis (35), systemic sclerosis (36), or rheumatoid arthritis (37). In these diseases, HB-EGF was mainly expressed by M1 macrophages as we observed in GCA. However, HB-EGF also appears to be expressed by M2 macrophages in post-inflammatory conditions such as idiopathic pulmonary fibrosis, contributing to the development of pulmonary fibrosis (38).

The effect of both AREG and HB-EGF on macrophages seems to be absent or weak in our study, using the production of pro-inflammatory cytokines or the activation of the NFκB pathway. This does not exclude the possibility of an autocrine effect of AREG and HB-EGF, but their role on macrophages remains controversial. In a murine model of glomerulonephritis, AREG was shown to enhance myeloid cell infiltration, M1 polarization of macrophages and cytokine secretion (21). However, other data suggested that AREG did not alter their own activation state or expression of pro-inflammatory cytokines (24).

In contrast, our data strongly support a role for AREG and HB-EGF on EGFR-expressing cells via in a paracrine manner, mainly by activating cell survival, growth and migration. We have demonstrated that AREG and HB-EGF strongly activated the MAPK pathway and showed a significant effect on both VSMC migration and proliferation. The role of AREG and HB-EGF on cell migration and proliferation may partly explain why diseases involving AREG and/or HB-EGF may be associated with cell proliferation and fibrosis (17,35,36,39), and many studies have demonstrated a role for both AREG and HB-EGF in organ fibrosis such as cirrhosis, pancreatic or lung fibrosis (40–42). AREG can stimulate the MAPK/ERK pathway in VSMC (43), through the activation of EGFR, and the key role of EGFR in VSMC migration and proliferation has been previously demonstrated (44,45). AREG have also been shown to be involved in promoting myofibroblast differentiation and fibrosis through the activation of the EGFR signaling pathway (31). Since one of the main mechanisms involved in GCA is the phenotypic change of VSMC into myofibroblasts, we could hypothesize that AREG is involved in VSMC differentiation into myofibroblasts and thus participate in vascular remodeling in GCA. Similarly, HB-EGF plays critical roles in proliferation, migration, differentiation, tissue repair, and regeneration and its role in cellular proliferation and migration appears to be even greater than that of AREG (46,47). HB-EGF is able to induce VSMC proliferation, but it also increases the secretion of IL-6 in VSMC, through the NF-κB pathway (44), data that we could not confirm. Overall, AREG and HB-EBF could show some redundant but also non redundant roles in inflammatory diseases, and especially GCA.

Our data raised the question of the potential interest of targeting AREG, HB-EGF or EGFR in GCA. Treatments targeting EGFR exist and are mainly used in malignancies, but EGFR inhibitors are associated with many side effects (48). Direct targeting of AREG and HB-EGF may be another option. In a mouse model of psoriasis, the use of anti-AREG antibodies in a SCID mouse transplanted with skin from psoriasis patients suppressed excessive skin proliferation (49). Other studies have also focused on anti-HB-EGF antibodies showing a potential interest of this treatment in mouse model of ovarian cancer (50). Targeting EGFR ligands may have fewer side effects than targeting EGFR itself. However, data on their efficacy and safety in inflammatory conditions are still lacking.

In conclusion, our study shows that the EGFR signaling pathway may play a role in GCA, especially AREG and HB-EGF. AREG and HB-EGF are expressed by infiltrating macrophages and induce EGFR-dependent activation of the MAPK pathway in VSMC, leading to proliferation and migration. Further studies are needed to explore the other EGFR ligands and to clarify their role in inflammatory conditions, especially in the dialog between macrophages and resident vascular cells.

## Authors’ contributions

KC and LD conducted experiments, acquired, and analyzed data. PB and MP helped for experiments. B. Terrier collected human TAB from Cochin institute and serums from VASCO cohort. B. Terris and PB provided temporal artery biopsies and helped for the interpretation of histologic data. KC and JD established the Opal panel. KC, LD, BT designed the study, analyzed data, and wrote the manuscript. Each author has proofread and approved the final version of the manuscript.

KC and LD contributed equally to this work.

## Acknowledgements

We thank all people who support the Fonds IMMUNOV.

We thank PARCC’s platforms: Corinne Lesaffre and Mathilde Lemitre from histological platform, Camille Knosp from cytometry platform and all the administration members.

## Ethics approval and consent to participate

Serum samples and TAB were obtained from patients enrolled in the VASCO (VASculitis COhort) study, a prospective cohort of patients with systemic vasculitides (NCT04413331), which includes a biological collection approved by the Direction de la Recherche Clinique et Innovation (DRCI) and the Ministère de la Recherche (N°2019-3677).

## Consent for publication

Not applicable

## Availability of data and materials

The datasets used and/or analysed during the current study are available from the corresponding author on reasonable request.

## List of abbreviations

α-SMA: α-smooth muscle actin
AREG: amphiregulin
EGF: Epidermal growth factor
EGFR: Epidermal growth factor receptor
ERK: Extracellular signal-regulated kinase
FBS: Fetal bovine serum
GCs: Glucocorticoids
GCA: Giant cell arteritis
GSEA: gene set enrichment analysis
HAoSMC: Human aorta, smooth muscle cells
HB-EGF: Heparin-binding epidermal growth factor
IL-6: Interleukin-6
IL-17: Interleukin-17
LPS: Lipopolysaccharide
MAPK: Mitogen-activated protein kinases
PBS: Phosphate buffer saline
VSMC: Vascular smooth muscle cells
TAB: Temporal artery biopsy
TNF-α: Tumor Necrosis Factor-α

